# Hidden hearing loss in a Charcot-Marie-Tooth type 1A mouse model

**DOI:** 10.1101/2023.12.14.571732

**Authors:** Luis R. Cassinotti, Lingchao Ji, M. Caroline Yuk, Aditi S. Desai, Nathan D. Cass, Zahara A. Amir, Gabriel Corfas

**Author notes:** Corresponding author: Gabriel Corfas, Ph.D. Kresge Hearing Research Institute The University of Michigan, Medical Sciences I Building, Rm. 5428 1150 West Medical Center Drive, Ann Arbor, MI 48109-5616. Lingchao Ji: Dept. of Otolaryngology, Peking University Shenzhen Hospital, China. Aditi S. Desai: University of Virginia School of Medicine, Charlottesville, Virginia, United States. Nathan D. Cass: Department of Otolaryngology, University of Kentucky, Lexington, KY, United States. **Conflict of Interest Statement:** GC was a scientific founder of Decibel Therapeutics, had equity interest in the company and received compensation for consulting. The company partially supported this work.

## Abstract

Hidden hearing loss (HHL), a recently described auditory neuropathy characterized by normal audiometric thresholds but reduced sound-evoked cochlear compound action potentials, has been proposed to contribute to hearing difficulty in noisy environments in people with normal hearing thresholds, a widespread complaint. While most studies on HHL pathogenesis have focused on inner hair cell (IHC) synaptopathy, we recently showed that transient auditory nerve (AN) demyelination also causes HHL in mice. To test the impact of myelinopathy on hearing in a clinically relevant model, we studied a mouse model of Charcot-Marie-Tooth type 1A (CMT1A), the most prevalent hereditary peripheral neuropathy in humans. CMT1A mice exhibited the functional hallmarks of HHL together with disorganization of AN heminodes near the IHCs with minor loss of AN fibers. These results support the hypothesis that mild disruptions of AN myelination can cause HHL, and that heminodal defects contribute to the alterations in the sound-evoked cochlear compound action potentials seen in this mouse model. Also, these findings suggest that patients with CMT1A or other mild peripheral neuropathies are likely to suffer from HHL. Furthermore, these results suggest that studies of hearing in CMT1A patients might help develop robust clinical tests for HHL, which are currently lacking.

## Introduction

Many people report having hearing difficulty in noisy environments despite presenting with normal audiometric thresholds (1–5). There is substantial evidence associating this auditory pathology with aging or with a history of “mild” noise exposures that only induce temporary increases in hearing thresholds (6–13). Studies on animal models demonstrated that such mild noise-exposures do not result in hair cell (HC) loss but instead destroy a subset of synapses between inner hair cells (IHCs) and spiral ganglion neurons (SGNs) (14). Moreover, in animals, IHC synaptopathy precedes HC loss during aging, even in the absence of noise over-exposure (14–20). Physiological recordings show that animals with IHC synaptopathy have normal hearing thresholds but reduced amplitudes of sound-evoked cochlear compound action potentials (15, 16, 18, 21), a condition that has been termed ‘hidden hearing loss’ (HHL) (22–24). Importantly, we recently demonstrated that young mice with inner hair cell synaptopathy independent of noise exposure have defects in temporal auditory processing (25), and histopathological studies have shown that age-related synaptopathy also occurs in humans (26, 27), suggesting that HHL might contribute to the hearing difficulties in people with normal thresholds. The compelling information generated from noise-exposed and aging animals led to the sense that IHC synaptopathy might be the only cause of HHL. However, more recently, we showed that transient auditory nerve (AN) demyelination causes a similar hearing deficit in mice without affecting IHC synapses (28), suggesting that demyelinating peripheral neuropathies could also cause HHL. To begin to test this possibility in clinically relevant experimental models, we focused on mouse models of Charcot-Marie-Tooth type 1 (CMT1) disease, the most common hereditary peripheral neuropathy in humans.

CMT1 is a type of CMT, a group of hereditary peripheral neuropathies that are associated with a range of genetic lesions and pathogenic mechanisms (29, 30). CMT1, which is considered a demyelinating CMT, includes multiple subtypes (from A to E) arising from alterations in diverse myelin-related genes (31). CMT1A, the predominant subtype (>50% of CMT1 cases), is caused by the duplication of *PMP22*, a gene that encodes an integral membrane glycoprotein essential for the formation and maintenance of compact myelin (32, 33). Interestingly, recent studies suggest that CMT1A patients might suffer from hearing difficulties consistent with HHL (34–37). However, this has not been confirmed due to the lack of validated clinical HHL tests in humans (23, 38). In contrast, patients with CMT1E, which is caused by a wide variety of *PMP22* point mutations (31) and lead to more severe neuropathy phenotypes with earlier onset, suffer from overt hearing loss (elevated auditory thresholds) (39–43). Furthermore, a CMT1E mouse model, Trembler-J, has been shown to have profound deafness including AN fiber loss (44, 45).

Here we used CMT1A (46) and CMT1E (a.k.a. Trembler-J) (47) mouse models to explore the specific impact of these hereditary peripheral neuropathies on inner ear structure and function. Our functional studies demonstrate that, as shown by others (44, 45), CMT1E mice present with early-onset overt hearing loss with profound auditory threshold shifts. In contrast, CMT1A mice exhibit the distinct electrophysiological features of HHL, i.e., normal auditory thresholds but reduced sound-evoked compound action potential amplitudes. However, in contrast to mice with IHC synaptopathy, sound-evoked compound action potential latencies are longer in CMT1A mice. At the structural level, whereas CMT1E present severe loss of AN fibers, the cochlear defects of CMT1A mice are primarily disorganization of AN heminodes near the IHCs and a subtle but significant miss localization of some nodes of Ranvier (NoR) to the area close to the heminodes.

Together, these results support the hypothesis that mild disruptions of AN myelination can cause HHL, and that AN heminodal defects are responsible for the reduced amplitude and longer latencies of sound-evoked compound action potentials seen in the mouse models (28, 48). Our results also support the notion that CMT1A patients are likely to suffer from HHL, and that this type of hearing disorder might affect patients with other types of peripheral neuropathies, including Guillain-Barré Syndrome. Based on these findings, we propose that studies of hearing in CMT1A patients might help in the design and validation of robust clinical tests for HHL. This would circumvent the difficulties encountered in the attempts to develop HHL tests based on self-reporting on historical noise exposure, which have resulted in conflicting conclusions (1, 38, 49).

## Results

### CMT1A mice have progressive hidden hearing loss

To determine the auditory phenotypes of CMT1A and CMT1E mice, we recorded distortion product otoacoustic emissions (DPOAEs) and auditory brainstem responses (ABRs) at 1, 2, 3 and 4 months of age (Figure 1A) in the mutants and their wildtype C57BL/6J littermate controls. ABR waveforms (Figure 1B) reflect the sound-evoked electrical activity along the ascending auditory pathway, from the activation of IHCs and SGNs in the cochlea up to the inferior colliculus in the central nervous system. ABR thresholds provide information about hearing sensitivity whereas the amplitude of the first peak of the ABR waveform reflects the magnitude of the sound-evoked SGN activation. DPOAEs are sounds generated by the inner ear in response to sound stimulation, reflecting the activity of outer hair cells (OHCs). Together, these measurements provide insights into the integrity of the cochlea. For example, increased ABR thresholds without a change in DPOAEs provides strong evidence for auditory neuropathy, i.e., defects in the function of IHCs and/or SGNs without OHC defects.

**Figure 1.**
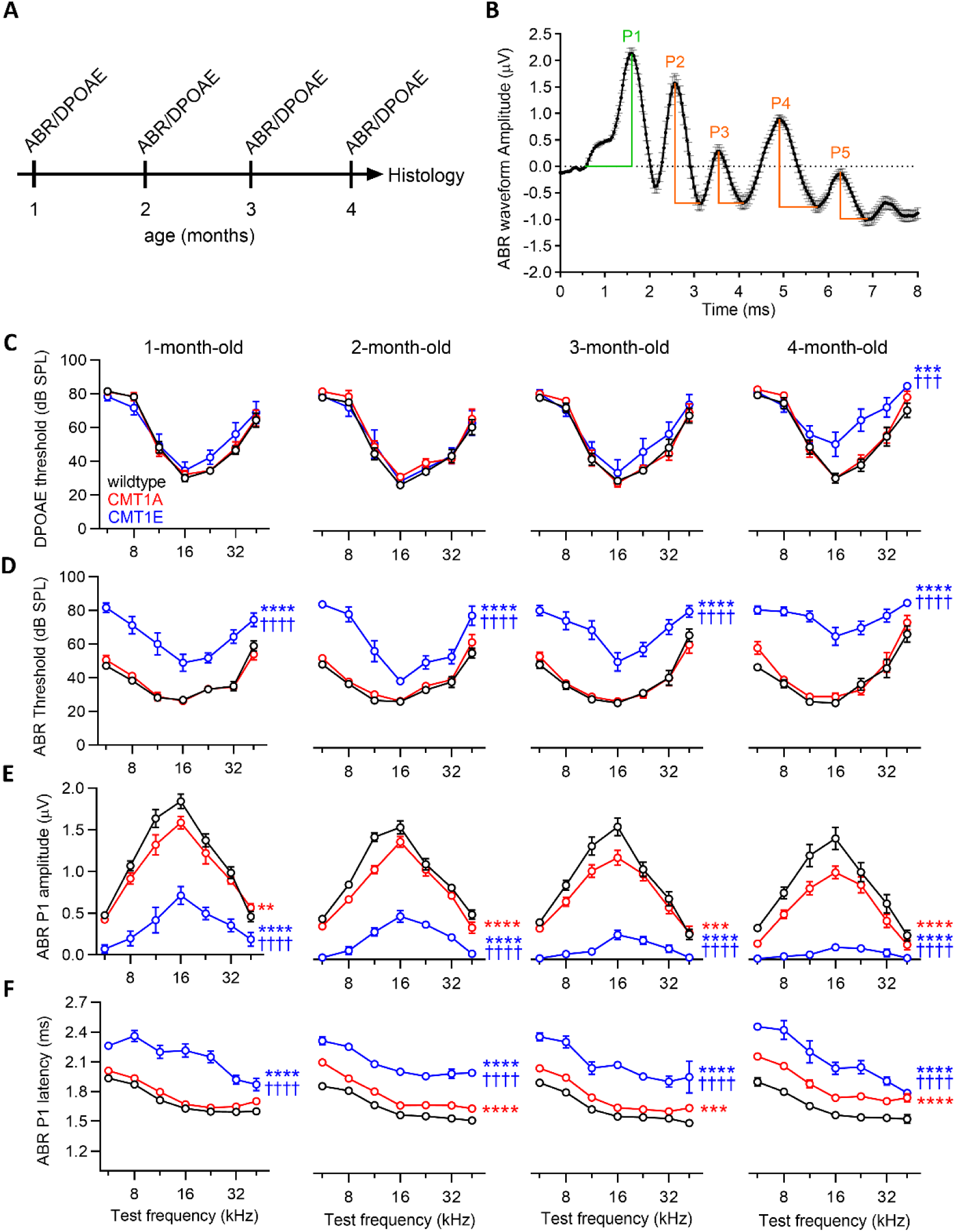
CMT1A mice have progressive hidden hearing loss whereas CMT1E have early onset overt hearing loss. **(A)** Experimental design. CMT1A, CMT1E and wildtype C57BL/6J littermates were used. ABRs and DPOAEs were recorded at 1, 2, 3 and 4 months of age and after the last tests, cochleae were collected and processed for either transmitted electron microscopy or confocal microscopy to evaluate structural changes at the level of the peripheral auditory nerve. **(B)** ABR waveform illustrating the criteria used to measure ABR peaks. Peak I amplitudes are measured relative to the baseline while peak II to peak V are measured from the top to the bottom right of each peak, respectively. **(C)** CMT1A mice have normal DPOAE thresholds at all time points while CMT1E mice show mild DPOAE threshold shifts starting at 4 months of age. **(D)** CMT1A mice have normal ABR thresholds while in CMT1E animals they are increased at all time points when compared with wildtype littermates. **(E)** ABR peak I amplitudes are progressively reduced in both CMT1 groups compared with wildtype mice; however, these reductions are more severe in CMT1E mice (also see Supplemental Figure 1). **(F)** ABR peak I latencies are longer in both CMT1 groups compared with wildtype mice, yet CMT1E animals show a more severe phenotype compared with CMT1A mice. Wildtype n = 14–19 mice; CMT1A n = 8–11 mice; CMT1E n = 6–12 mice. DPOAE threshold, ABR threshold, ABR peak I amplitude or ABR peak I latency at each individual time point were evaluated by two-way ANOVA followed by Tukey’s multiple comparisons test. ABR peak I amplitudes and latencies were measured at 80 dB SPL. ** p< 0.01; *** p< 0.001; **** p< 0.0001 vs wildtype mice; ††† p< 0.001; †††† p< 0.0001 vs CMT1A mice. Error bars represent SEM.

As anticipated for C57BL/6J wildtypes, control mice show stable DPOAE and ABR thresholds during their first 4 months of life (Figure, 1C-D), and a small but statistically significant age-related decline in ABR peak I amplitudes (Figure 1E; Figure 2A, Supplemental Figure 1 and Supplemental Figure 2A). Their ABR peak I latencies become shorter by 2 months of age, possibly due to the maturation of AN myelination (50, 51) (Figure 1F and Supplemental Figure2B). Consistent with published reports (44, 45), CMT1E mice have higher ABR thresholds at all ages and frequencies tested (Figure 1D) while their DPOAE thresholds are mildly increased compared to wildtype mice only at 4 months of age (Figure 1C). Furthermore, CMT1E ABR peak I amplitudes are considerably lower and the latencies longer than in wildtype mice (Figure 1, E-F; Figure 2A and Supplemental Figure 1). These findings support a profound auditory neuropathy as early as 1 month of age in CMT1E mice, which then progresses to include OHC dysfunction. In contrast, CMT1A mice have normal ABR and DPOAE thresholds at all ages (Figure 1, C-D), but their ABR peak I amplitudes are already smaller than those in wildtypes by 1 month of age and decline further over time (Figure 1E; Figure 2A and Supplemental Figure 1), the hallmarks of progressive HHL. Moreover, ABR peak I latencies are longer than those in wildtype mice at 2 month of age and do not get shorter at later time points (Figure 1F and Figure 2A), suggesting a lack of myelin maturation. The ABR waveforms (Figure 2A) also show that the summating potentials (SP), which represent the activation of IHCs, are normal in CMT1A mice but are significantly reduced in 4-month-old CMT1E, likely reflecting the OHC dysfunction seen at this age. The waveforms also illustrate that CMT1E mice have a progressive decrease in signaling along the ascending auditory pathway (Figure 2), whereas ABR peak II, III, IV and V in CMT1A mice are normal at 4 months of age (Figure 2), suggestive of the homeostatic compensation in the auditory brainstem and midbrain seen in animals with partial peripheral deafferentation (52–56). Remarkably, after 2 months of age, the functional defects in CMT1A are identical to those we previously found in mice following transient AN demyelination, i.e., normal threshold, reduced ABR peak I amplitudes, and longer peak I latencies (28).

**Figure 2.**
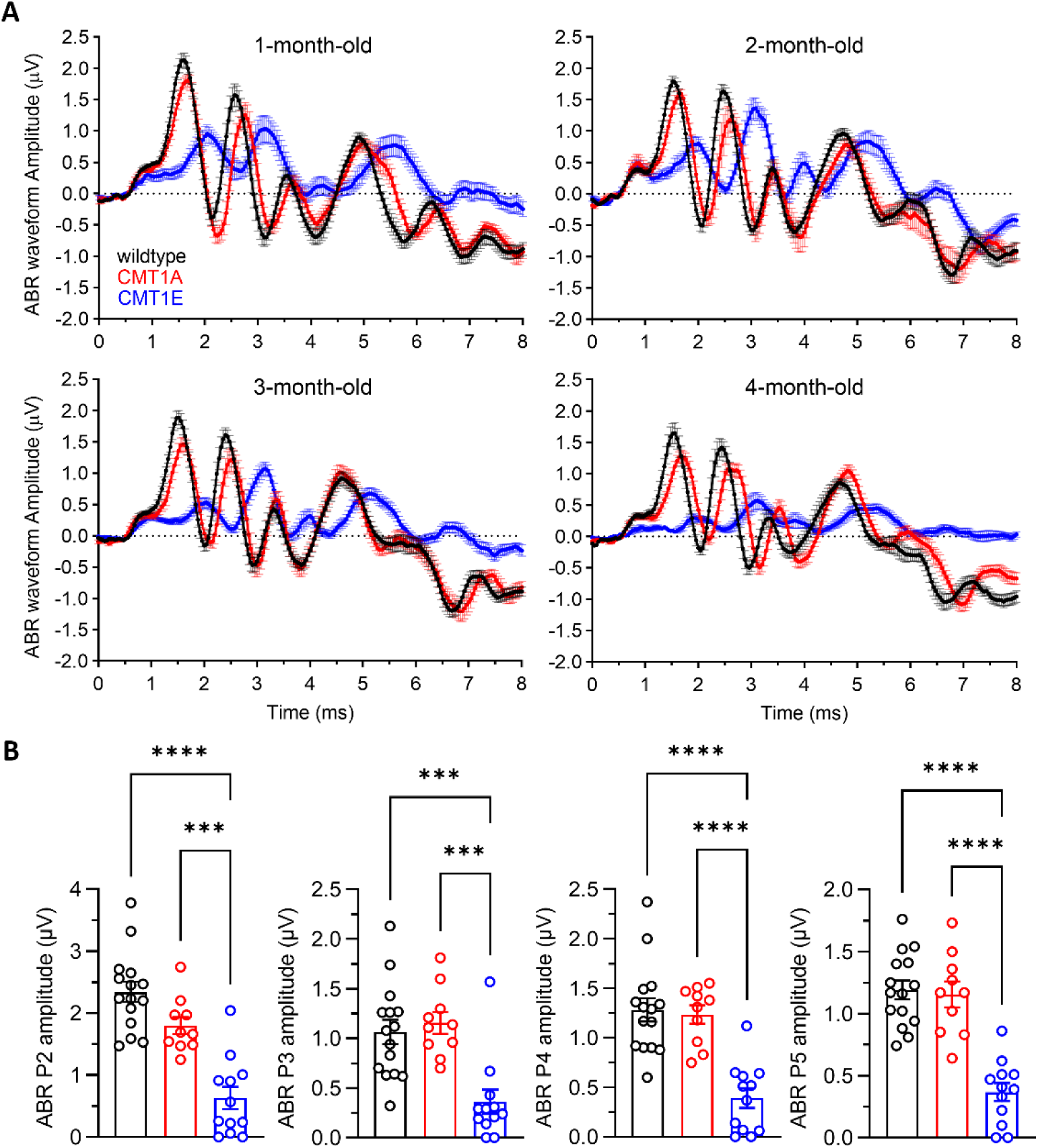
Distinct effects of CMT1A and CMT1E on signaling along the ascending auditory pathway. **(A)** Mean ABR waveforms recorded at 1, 2, 3 and 4 months of age show a progressive delay and reduction of the first ABR peak in both CMT1 mouse models. CMT1E phenotype is more severe than in CMT1A mice, affecting the later ABR peaks along the ascending auditory pathway. ABRs shown here are group means in response to 16 kHz tone pips at 80 dB SPL. Error bars represent SEM. **(B)** Quantification of ABR amplitudes in peak II, III, IV and V at 4 months of age shows that CMT1E mice have decreased signaling along the ascending auditory pathway while in CMT1A mice are normal compared with wildtype mice. ABR peak II-V amplitudes were evaluated at 16kHz cochlear frequency and at suprathreshold levels (80 dB SPL). Wildtype n = 15 mice; CMT1A n = 10 mice; CMT1E n = 12 mice. One-way ANOVA followed by Tukey’s multiple comparisons test was used to evaluate statistical differences among the experimental groups. *** p < 0.001; **** p < 0.0001. Error bars represent SEM.

### CMT1A and CMT1E mice have distinct inner ear axonal and myelin pathologies

To explore the structural basis for the functional phenotypes in the CMT1 mice, inner ears were harvested after the final ABR/DPOAE recording and subjected to several levels of histological analysis. Transmission electron microscopy of cross sections through the osseous spiral lamina (OSL, the bony structure encasing the AN fibers, Supplemental Figure 3) in the mid-cochlear region (∼16 kHz) demonstrated that CMT1A cochleas have a small but significant reduction in myelinated axon density (18 %) whereas axonal loss is much larger in CMT1E mice (∼78 %) (Figure 3, A-B, Supplemental Figure 3 and Supplemental Figure 4). Interestingly, while the diameters of the remaining axons in CMT1E cochleas are like those seen in wildtypes, axonal diameters in CMT1A ears are reduced (Figure 3C). In contrast, measurements of g-ratio (the ratio of the inner versus outer layer diameter of the myelin sheath), reflect myelin is thinner (larger g-ratio) in CMT1E auditory axons (Figure 3, D and E), whereas CMT1A cochleas have a more complex myelin phenotype, with small diameter axons exhibiting thicker myelin, resulting in a steepening of the regression line in the g-ratio vs axon diameter graph (Figure 3, D and E). In addition, in CMT1A we observed some axons with mild myelin compaction defects, but myelin abnormalities were more dramatic in CMT1E mice (Supplemental Figure 4). These results demonstrate that CMT1A and CMT1E mutations cause distinct AN axonal and myelin structural changes which are similar to those seen in other peripheral nerves (46, 57–62), with CMT1A mice presenting a less severe phenotype.

**Figure 3.**
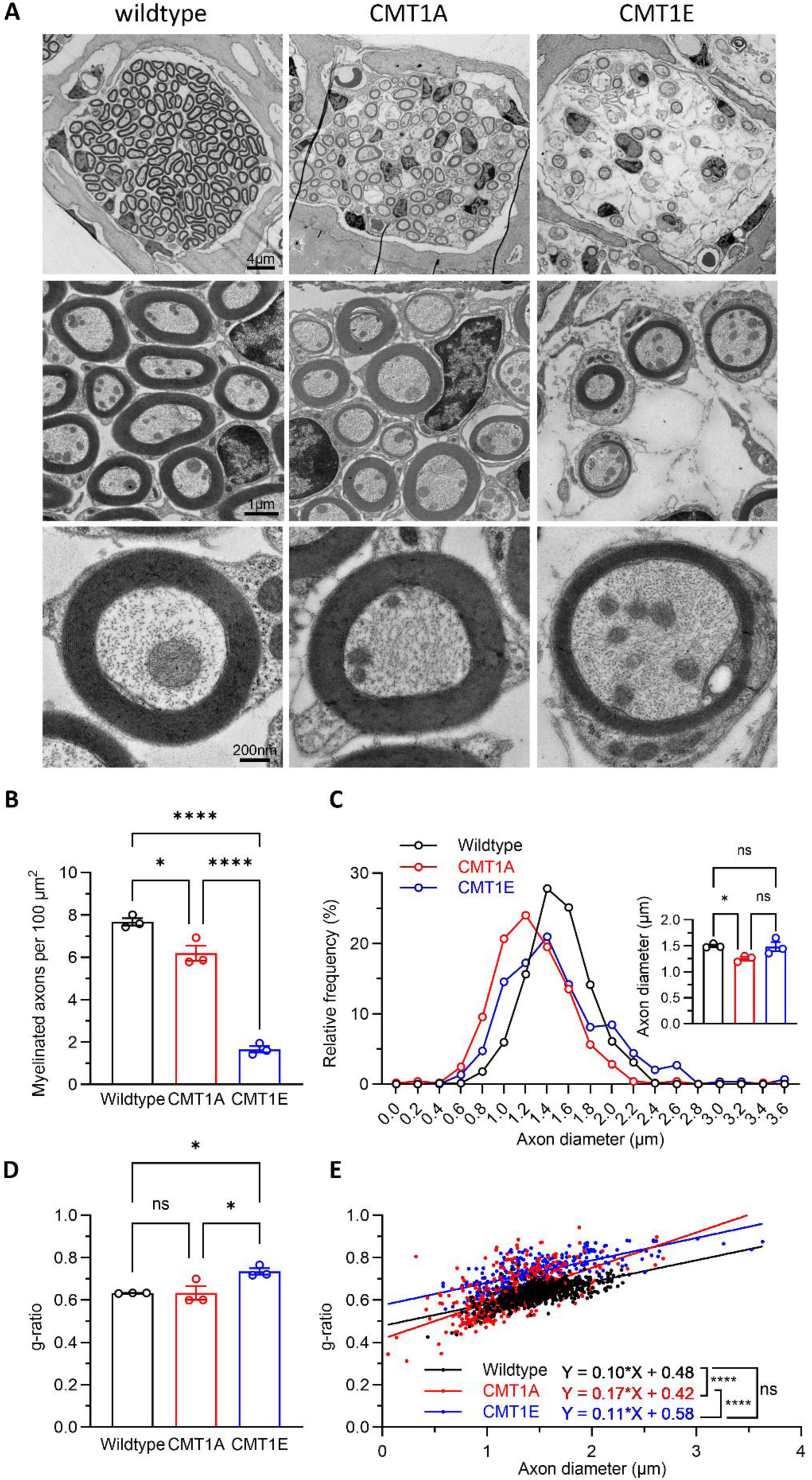
CMT1A and CMT1E mice have distinct myelin and axonal pathologies. **(A)** Representative electron micrographs of sections through the OSL showing an axon bundle (top), a high magnification image of the axons (middle) and an individual myelinated axon image (bottom) from 4-month-old wildtype (left), CMT1A (center) and CMT1E (right) mice. **(B)** CMT1A and CMT1E animals have different degrees of myelinated axons loss, 19 % and 78 % respectively compared to wildtype mice. Dots represent the number of myelinated axons per 100 µm^2^ cross section through the OSL at 16 kHz per mouse (n = 3 cochleae per genotype). **(C)** Frequency histogram showing that CMT1A mice have a larger number of small diameter axons than wildtype mice. Inset: CMT1A mice have a 16% reduction in axon diameter vs wildtype. Each dot represents the averaged axon diameter per experimental animal (n = 3 cochleae per genotype). **(D)** CMT1E mice have decreased myelin thickness (g-ratio = 0.735) compared with wildtype (g-ratio = 0.633) and CMT1A mice (g-ratio = 0.633). Each dot represents the averaged g-ratio per experimental animal (n = 3 cochleae per genotype) **(E)** CMT1E hypomyelination is independent of axon diameter while CMT1A mice have an increased steepening in the regression line in the g-ratio vs axon diameter graph (n= 3 cochleae per genotype at the 16 kHz region; wildtype axons = 201-253 per cochlea, CMT1A axons = 138-198 per cochlea, CMT1E axons = 62-146 per cochlea). Quantification of myelinated axon densities, axon diameters and g-ratios were analyzed by one-way ANOVA followed by Tukey’s multiple comparisons test. Linear regression and differences among slopes of axon diameters vs g-ratio plots were obtained with GraphPad Prism simple linear regression test. * p < 0.05; ** p < 0.01; **** p < 0.0001. Error bars represent SEM.

### CMT1A results in AN heminodes abnormalities

The auditory physiology phenotypes in CMT1A mice, i.e., normal threshold, reduced ABR peak I amplitudes, and longer peak I latencies (Figure 1, Figure 2A, and Supplemental Figure 1), are remarkably similar to those we previously recorded in mice that had undergone remyelination after transient ablation of Schwann cells (28). In the latter case, the only AN structural abnormality that correlates with the physiological findings is the disarray of AN heminodes, the nodal structures at the peripheral ends of the terminal Schwann cells adjacent to the IHCs. AN heminodes have been proposed to function as the SGN action potential initiation site, and to play a critical role in synchronous neural transmission (48, 63–65). To evaluate the impact of CMT1A and CMT1E on nodal structures, we used laser confocal microscopy with antibodies targeted to Ankyrin-G (a nodal protein marker (66)), Caspr (a paranodal marker (66)), and Myelin Basic Protein (MBP) to visualize myelin (67) on cochleas from 4-month-old mice.

Whereas AN heminodes in wildtype mice are tightly clustered within the 20 µm segments of the myelinated axon closest to the IHCs, heminode clustering is disrupted in CMT1A cochleas (Figure 4, A and B), with elongated Caspr-positive regions and Ankyrin-G-positive regions located further from the peripheral end of the heminodes relative to those in wildtypes. Furthermore, similar to what we observed following transient demyelination-remyelination (28), there are numerous nodes of Ranvier (NoR) in the area directly adjacent to heminodes in CMT1A cochleas, whereas very few NoR are present in this area in wildtypes (Figure 4, B and D).

**Figure 4.**
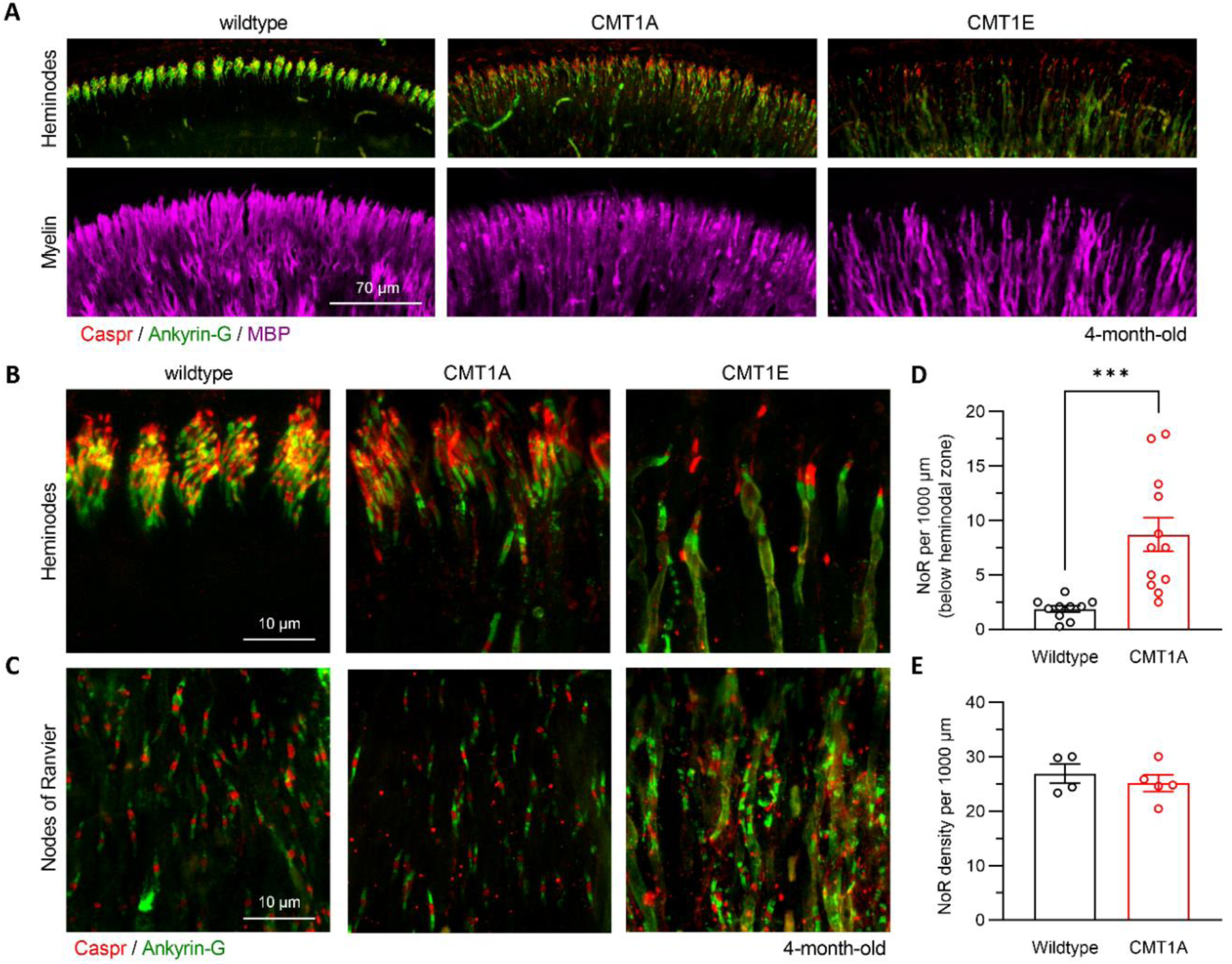
CMT1A and CMT1E mice have nodes of Ranvier and heminodes abnormalities. **(A)** Representative low magnification images showing that the AN heminodes adjacent to the organ of Corti (top) and the AN myelin in the OSL (bottom) are disrupted in CMT1A (center) and CMT1E (right) cochleae at 4 months old compared with wildtype mice (left). **(B)** High magnification images showing the heminodes and nodes of Ranvier and **(C)** the nodes of Ranvier at the 16 kHz cochlear region from wildtype (left), CMT1A (center) and CMT1E (right) mice. **(D)** CMT1A mice have increased number of nodes of Ranvier-like structures in the 20 µm area below the disrupted heminodes compared to wildtypes (wildtype n = 10 cochleae; CMT1A n = 12 cochleae). **(E)** The density of nodes of Ranvier at the level of the OSL in CMT1A mice is not affected. Samples were immunolabelled for paranodes (Caspr); nodes of Ranvier (Ankyrin-G) and myelin (MBP). Wildtype n = 4 cochleae; CMT1A n = 5 cochleae. One cochlea from each mouse was analyzed and quantifications were done at the ∼16 kHz region. Nodes of Ranvier density in either the 20µm below the heminodal area or in the OSL region was analyzed by two-tailed t-test. Quantification of heminodes and nodes of Ranvier in CMT1E mice was not possible due to the severe axonal damage exhibited. *** p < 0.001. Error bars represent SEM.

Nevertheless, the overall NoR density in the peripheral AN processes of CMT1A mice is similar to that in wildtypes (Figure 4, C and E), suggesting that, overall, Schwann cell internode lengths are not affected in these mutants. As can be expected from the severe AN axonal loss in CMT1E mice, the heminodal disruption in these mutants is more severe (Figure 4, A and B). Accurate quantification of heminodes and NoR density in CMT1E mice was not possible due to the profound AN axon loss.

### CMT1A and CMT1E mice have different degrees of IHC-SGN synapse loss without affecting hair cell number

Finally, to evaluate the potential contribution of HC or IHC synapse loss to the auditory physiology defects of the mutant mice, we quantified the number of IHCs and OHCs as well as IHC-SGN synaptic density in 4-month-old mice from each cohort. Notably, IHC and OHC survival were not affected by either of the two CMT1 mutations (Figure 5, A and C). In concordance with the severe axonal loss (Figure 3, A and B), CMT1E mutants have a large (58–61 %) reduction in IHC-SGN synaptic density compared with wildtypes (Figure 5, B and D). In contrast, CMT1A mice exhibit a more modest (15 %) reduction in IHC-SGN synaptic density in the mid-high frequency area (Figure 5, B and D), which may also reflect the small degree of axonal loss in these mutants (Figure 3, A-B, Supplemental Figure 3, and Supplemental Figure 4).

**Figure 5.**
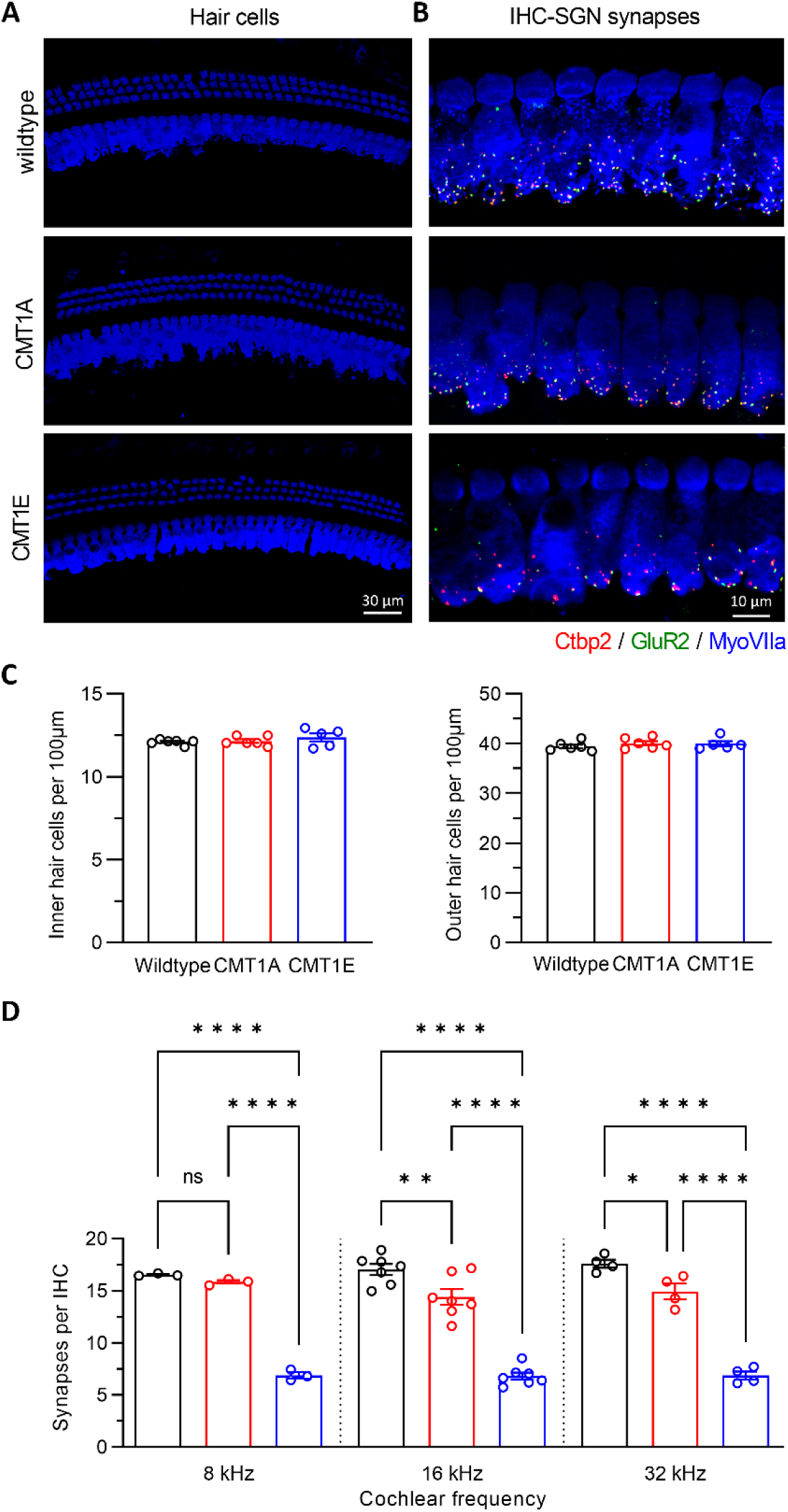
CMT1A and CMT1E mice have different degrees of IHC-SGN synapse loss without hair cell loss. Representative confocal images of hair cells **(A)** and IHC-SGN synapses **(B)** at the 16 kHz region of wildtype (top), CMT1A (middle) and CMT1E (bottom) cochleae immunolabeled for hair cells (MyoVIIa), pre-synaptic ribbons (Ctbp2) and post-synaptic receptor patches (GluR2). **(C)** Inner (left) and outer (right) hair cells survival is not affected by these two different CMT1 mutations. Wildtype n = 6 cochleae; CMT1A n = 6 cochleae; CMT1E n = 5 cochleae. One cochlea from each animal was imaged, with 3 adjacent z-stacks acquired at each specific cochlear region. **(D)** CMT1A mice have mild IHC-SGN synapse loss at 16 and 32 kHz areas, compared with wildtype mice. CMT1E IHC-SGN synapse density is more severely reduced than in CMT1A. Wildtype n = 3-7 cochleae; CMT1A n = 3-7 cochleae; CMT1E n = 3-7 cochleae. One cochlea from each animal was imaged, with 3 adjacent z-stacks acquired at each specific cochlear region. IHC-SGNs synapse density was analyzed by one-way ANOVA followed by Sidak’s multiple comparisons test. Quantification of inner and outer hair cells was analyzed by Kruskal-Wallis test followed by Dunn’s multiple comparisons test. * p < 0.05; ** p < 0.01; **** p < 0.0001. Error bars represent SEM.

## Discussion

The concept of HHL was first proposed in 2011 (24), and since then research on the mechanism of its pathogenesis has centered around IHC synaptopathy (14). Here we provide evidence that a mouse model of CMT1A, the most prevalent hereditary peripheral myelin disorder, exhibits the functional hallmarks of HHL, i.e., normal auditory thresholds but reduced ABR peak I amplitudes. The observation that a mouse model of CMT1E, which is caused by different alterations in the same gene as CMT1A (*PMP22*), has overt hearing loss (like some CMT1E patients) together with a distinct set of structural deficits in the cochlea, corroborates the specificity of the CMT1A phenotype. Furthermore, the notion that myelin disorders are an alternative etiology for HHL is supported by the remarkable phenotypic similarities between mice following transient AN demyelination (28) and the CMT1A model, i.e., normal ABR and DPOAE thresholds, reduced ABR peak I amplitudes together with longer peak I latencies. This idea is also supported by the fact that the longer ABR peak I latencies observed in both myelinopathy models is absent from mice with noise-induced or age-related IHC synaptopathy (14, 28).

Here we also show that, like in mice following transient AN demyelination (28), cochlear heminodes are disrupted and some NoR are mislocalized in CMT1A mice. Moreover, this structural phenotype correlates with the reduced sound evoked SGN activation in both models, supporting the crucial role of the AN heminodes as the SGN action potential initiation site, and its importance for fast and synchronized AN spike generation as has been previously suggested in computational models or electrophysiological and immunostaining studies (48, 63–65). Importantly, our mathematical model of HHL caused by myelinopathy suggest that the heminodal defects might cause AN fiber spike generation failure which could contribute to the reduced amplitude of the first peak of the ABR waveform (48, 63–65). The finding that CMT1A results in AN heminode defects also raises the possibility that defects in other heminodes; e.g., those close to neuromuscular synapses, or other action potential initiation sites, e.g., axon initial segments in sensory neurons, might contribute to other sensory and motor phenotypes in CMT1A.

The distinct AN myelin and axonal phenotypes in each CMT1 mouse model are also remarkably similar to those observed in other peripheral nerves, suggesting that the specific *PMP22* mutations exert common pathological effects across myelinating glia. For example, AN in CMT1A cochleas exhibit an increased proportion of smaller axons, a slight reduction in the number of myelinated fibers, hypermyelination of small diameter fibers, and a steepening of the regression line in the g-ratio vs axon diameter graph, all features that have been reported in other peripheral nerves of this mouse model and also in CMT1A rat models (46, 57, 58, 60). In contrast, sciatic nerves from CMT1E mice exhibit hypomyelination, much like what we and others have observed in the cochlea of this model (59, 61, 62). Furthermore, the AN phenotypes in both models are consistent with findings in other nerves in CMT1 patients (68).

Given the lack of validated diagnostic tools for HHL in humans (23), our findings in the CMT1A mouse model provide strong support for the handful of clinical observations suggesting CMT1A patients have HHL-like phenotypes. An early study of children with CMT1A reported that most have normal audiological thresholds, but some have lower ABR amplitudes and longer latencies (36). Furthermore, more than 60% of these children have defects in auditory perception, temporal processing, and speech recognition, consequences that have now been predicted for humans with HHL (1–9). More recent studies reported that adult CMT1A patients have normal auditory thresholds, but defects in speech perception in noisy backgrounds, reflected by lower temporal and spectral resolution than healthy controls (34, 35). Importantly, our results also suggest that studying patients with genetic disorders such as CMT1A is likely to facilitate the development of better tests for HHL that could be validated using the power of phenotype-genotype relationships.

The auditory function deficits of CMT1A and CMT1E mice are profoundly different, but in each case are similar to those seen in patients with the respective mutations (34–37, 39–43, 69). Thus, we infer that it is likely that the cochlear structural phenotypes in each patient population correlate to those we see in each mouse model as well. Specifically, our findings suggest that CMT1E patients most likely have severe SGN neuron loss, whereas CMT1A patients are likely to retain most SGNs but have heminode disruptions. If this is the case, it would imply that CMT1E patients are unlikely to benefit much from hearing aids or cochlear implants, whereas CMT1A patients might be helped by hearing augmentation. This could in part explain the different efficacy outcomes observed in some CMT cochlear implanted patients (70, 71).Given the growing evidence that hearing loss can contribute to cognitive decline and dementia (72–74), it would be also important that neurologists who care for peripheral neuropathy patients include auditory testing in clinical workups and consider hearing enhancement devices in treatment. Furthermore, our observations that a mouse model of myelinopathy has the hallmarks of HHL suggest that this might be a frequent comorbidity in other peripheral demyelinating neuropathies, e.g., Guillain-Barre Syndrome. In this context, therapies that promote AN remyelination and heminode reorganization could help to prevent or slow the progression of hearing impairments associated with CMT1A or other peripheral demyelinating neuropathies (75–79).

## Materials and Methods

### Sex as a biological variable

Our study examined male and female mice, and no differences were found between both sexes.

### Animals

*B6.Cg-Tg (PMP22) C3Fbas/J* mice (a.k.a. PMP22-C3 or C3-PMP, here termed CMT1A; The Jackson Laboratory, stock #030052), *B6.D2-pmp22Tr-j/J* mice (a.k.a. Trembler-J, here termed CMT1E, The Jackson Laboratory, stock #002504) and C57BL/6J wild type littermates (1 to 4 months old, male and female) were used in this study. CMT1A mice express 3 copies of wildtype human peripheral myelin protein 22 (PMP22) gene, mimicking CMT1A (46). CMT1E mice have a semi-dominant point spontaneous mutation, i.e., a T to C transition at nucleotide position 47, resulting in the substitution of a leucine by a proline in the first transmembrane domain of the PMP22 protein (47). This point mutation causes peripheral demyelination that mimics CMT1E disease.

### Auditory function tests

ABRs and DPOAEs were performed on mice anesthetized with a mixture of ketamine (100 mg/kg, i.p.) and xylazine (20 mg/kg, i.p.). All groups of animals were tested at 1, 2, 3 and 4 months of age.

For ABR recordings, acoustic stimuli were delivered through a closed acoustic system, consisting of two sound sources (CDMG15008-03A, CUI) and an electret condenser microphone (FG-23329-PO7, Knowles) as an in-dwelling probe microphone. Three needle-electrodes were placed into the skin at the dorsal-midline: one close to the neural crest, one behind the left pinna and one at the base of the tail (ground). ABR potentials were evoked with 5 ms tone pips (0.5 ms rise–fall, with a cos^2^ envelope, at 40 s^-1^) delivered to the eardrum at log-spaced frequencies from 5.6 kHz to 42.25 kHz. The response was amplified (10,000X) and filtered (0.3–3 kHz) with an analog-to-digital board in a PC-based data-acquisition system. The sound level was raised in 5 dB steps from 20 to 80 dB SPL. At each level, 1024 responses were averaged (with stimulus polarity alternated) after “artefact rejection” above 15 µV.

DPOAEs, in response to two primary tones of frequency f1 and f2, were recorded at (2 x f1) - f2, with f2/f1 = 1.2, and the f2 level 10 dB lower than the f1 level. Stimuli were raised in 5 dB steps from 20 to 80 dB. The ear-canal sound pressure was amplified and digitally sampled at 4 µs intervals. DPOAE thresholds were defined as the lower SPL where (2f1-f2) - (2f1-f2Nse) > 0.

Both ABR and DPOAE recordings were performed using the EPL cochlear function test suite (Mass Eye and Ear, Boston, MA, USA). ABR thresholds, ABR peak I amplitudes and latencies, ABR waveforms and DPOAE thresholds were analyzed with ABR peak Analysis software (Mass Eye and Ear, Boston, MA, USA) and Microsoft Excel.

### Plastic sections and transmission electron microscopy

Otic capsules from 4-month-old CMT1A, CMT1E and wildtype mice were dissected and fixed in 1.25 % paraformaldehyde 2.5 % glutaraldehyde 100 mM cacodylate overnight, followed by osmification in 1 % osmium tetroxide for 45 min and decalcification in 5 % EDTA 1 % glutaraldehyde 0.1M PBS for 5 days. Cochleae were microdissected and whole mount tissues were gradually dehydrated in ethanol (70 %, 95 % and 100 % steps) and embedded in araldite resin. The embedded samples were degassed for 2 h and hardened at 60°C for 5 days. For myelinated axons density counts, semi-thin cross sections of AN fibers were made at the level of the OSL through the 16 kHz cochlear region. All myelinated fibers in the OSL cross sections were counted at 60X magnification. The number of fibers was then divided by the axonal bundle extent of OSL included in that section, to arrive at an estimate of the number of myelinated fibers per 100 µm^2^ of organ of Corti. For g-ratio and axon diameter estimations ultra-thin (70 nm) OSL cross-sections were prepared for transmission electron microscopy. Ultra-thin sections containing 16 kHz cochlear region were sequentially post-stained with 6 % w/v uranyl acetate and 4.4 % w/v lead citrate. Transmission electron microscopy was performed on JEOL 1400-plus electron microscope (JEOL USA, Peabody, MA). Multiple non-overlapping regions of the AN fibers cross-sections were imaged at 600X, 3000X and 10000X magnification.

The circumference of each axon and axon + myelin sheath were measured using ImageJ software g-ratio plugin (version 1.53c, NIH, MD) on 3000X magnification images. g-ratios were calculated as g-ratio = (axon area)/(axon area + myelin sheath area). Axon diameters were calculated using the same software. Axons with circularity (4 x π x area/perimeter^2^) under 0.6 were excluded from analysis. All electron microscopy images are representative of 3 individual mice per group.

### Immunostaining and confocal imaging

Inner ear tissues from 4-month-old CMT1A, CMT1E and wildtype mice were dissected and fixed in 4 % paraformaldehyde in 0.01M phosphate-buffered saline (PBS) for 2 h at room temperature, followed by decalcification in 5 % EDTA at 4 °C for 5 days. Then, cochlear tissues were microdissected and permeabilized by freeze–thawing in 30% sucrose in PBS. The microdissected tissues were incubated in blocking buffer containing 5 % normal horse serum and 0.3 % Triton X-100 in PBS for 1 h. Tissues were then incubated in primary antibodies (diluted in 1 % normal horse serum and 0.3 % Triton X-100 in PBS) at 37 °C overnight. The primary antibodies used in this study were as follows; for nodes of Ranvier, heminodes and myelin staining: anti-Caspr clone K65/35 (NeuroMab, Davis, CA; 1:1000), anti-Ankyrin G (gift of Paul Jenkins laboratory, Department of Pharmacology, University of Michigan, Ann Arbor, MI, USA; 1:500), anti-MBP (Millipore, Billerica, MA; 1:1000; catalog no. MAB386); for hair cells and IHC-SGN synapses: anti-Ctbp2 (BD Biosciences, San Jose, CA; 1:200; catalog no. 612044), anti-GluR2 (Millipore, Billerica, MA; 1:1000; catalog no. MAB397) and anti-MyoVIIa (Proteus Biosciences, Ramona, CA; 1:100; catalog no. 25-6790). Tissues were then incubated with appropriate Alexa Fluor-conjugated fluorescent secondary antibodies (Invitrogen, Carlsbad, CA; 1:1000 diluted in 1% normal horse serum and 0.3% Triton X-100 in PBS; for nodes of Ranvier, heminodes and myelin: AF488 IgG1 catalog no. A-21121; AF568 catalog no. A11036; AF647 catalog no. A-21247; for hair cells and IHC-SGN synapses: AF488 IgG2a catalog no. A-21131; AF568 IgG1 catalog no. A-21124; AF647 IgG catalog no. A-21244) for 1 h at room temperature. The tissues were mounted on microscope slides in ProLong Diamond Antifade Mountant (Thermo Fisher Scientific). All pieces of each cochlea were imaged at low power (10X magnification) to convert cochlear locations into frequency (tonotopic mapping) using a custom plug-in to ImageJ (1.53c NIH, MD) available at the website of the Eaton-Peabody Laboratories (EPL). Cochlear tissues from the ∼16 kHz region were used for further analyses unless stated different. Confocal z-stacks of cochlear tissues were taken using a Leica SP8 confocal microscope.

Nodes of Ranvier and heminodal structure z-stacks (0.3 µm step size) were taken under 63X (+ 4.61X optical zoom) or 40X magnification (for low magnification images) at the ∼16 kHz cochlear region (one z-stack per individual animal). ImageJ/Fiji software (version 1.53c, NIH, MD) was used for z-stack processing and quantification of nodes of Ranvier presence either in the 20 µm below the heminodal zone or in the OSL lamina region. For nodes of Ranvier quantification, the number of nodes of Ranvier in each sample was counted manually using ImageJ/Fiji software multi-point counter tool. For figures, one representative image was selected from amongst the 12–48 images.

Images for hair cell counts were taken under 40X magnification. For inner hair cell synapse counts, z-stacks (0.3 μm step size) were taken under 63X (+2.4X optical zoom) magnification spanning the entire IHC height to ensure all synapses were imaged. Imaging and analyses of cochlear hair cells and synapses were performed as previously described in (80). Briefly, ImageJ/Fiji software (version 1.53c, NIH, MD) was used for image processing and quantification. One cochlea from each animal was imaged for each experiment, with 3 adjacent z-stacks acquired at each specific cochlear region. For hair cell quantification, the number of inner and outer hair cells at specific cochlear regions in each animal was determined based on the MyoVIIa channel and counted manually using ImageJ/Fiji software multi-point counter tool. For synapse counts, CtBP2 and GluR2 puncta in each image stacks were also captured and counted manually using ImageJ/Fiji software multi-point counter tool. Synaptic counts of each z-stack were divided by the number of IHCs, which could be visualized by staining of MyoVIIa antibody. Each individual image usually contained 8–10 IHCs. For figures, one representative image was selected from amongst the 9–24 images from the specific frequency shown.

### Statistical Analysis

Graphics and statistical tests were performed using GraphPad Prism version 9.3.1 for Windows (GraphPad Software, www.graphpad.com). Data sets with normal distributions were analyzed with parametric tests whereas non-parametric tests were used for sets that did not conform to normality criteria. We used two-way ANOVA, followed by Tukey’s multiple comparisons test to compare either DPOAE thresholds, ABR thresholds, ABR peak I amplitudes or latencies at each time point (1, 2, 3 and 4 months) among CMT1A, CMT1E and wildtype mice (Figure 1, C-F; Supplemental Figure 1 and Supplemental Figure 2). One-way ANOVA followed by Tukey’s multiple comparisons test was used to evaluate statistical differences in ABR peak II, III, IV and V amplitudes (Figure 2A). Quantification of myelinated axon densities, quantification of axon diameters and g-ratios were analyzed by one-way ANOVA followed by Tukey’s multiple comparisons test (Figure 3, B-D). Simple linear regression of axon diameters vs g-ratios and statistical differences among slopes were obtained with GraphPad Prism (Figure 3E).

Quantification of nodes of Ranvier density in either the 20µm below the heminodal area or in the OSL region were analyzed by two-tailed t-test (Figure 4, D-E). Quantification of inner and outer hair cells was analyzed by Kruskal-Wallis test followed by Dunn’s multiple comparisons test (Figure 5C). Quantification of IHC-SGNs synapse density was analyzed by one-way ANOVA followed by Sidak’s multiple comparisons test (Figure 5D). All p < 0.05 were considered as statistically significant.

### Study approval

All animal procedures were approved by the Institutional Animal Care and Use Committee of the University of Michigan, and all experiments were performed in accordance with relevant guidelines and regulations.

## Data Availability

All data generated or analyzed during this study are included in the manuscript and data values in the supporting XLS file. Original electron microscopy and confocal images used for calculations, ABR and DPOAE recordings will be uploaded to Dryad.

## Author Contributions

LRC and GC designed the research studies. LRC, LJ, MCY, ASD and NDC conducted experiments. LRC, MCY, ASD and NDC acquired data. LRC, MCY, ASD, NDC, ZAA and GC analyzed data. LRC, LJ, MCY, ASD, NDC and GC wrote the manuscript.

## Acknowledgments

This work was supported by NIH R01DC018500 (GC) and a grant from Decibel Therapeutics. We thank Dr. Paul Jenkins for providing the Ankyrin G antibody and Dr. David Kohrman for helpful comments on the manuscript.

## Abbreviation list

ABR: Aditory Brainstem Response

DPOAE: Distortion Product Otoacoustic Emissions

CMT: Charcot-Marie-Tooth disease

CMT1A: Charcot-Marie-Tooth type 1A

CMT1E: Charcot-Marie-Tooth type 1E

HHL: Hidden Hearing Loss

AN: Auditory Nerve

HC: Hair Cell

IHC: Inner Hair Cell

OHC: Outer Hair Cell

PMP22: Peripheral Myelin Protein 22

SGN: Spiral Ganglion Neuron

MBP: Myelin Binding Protein

NoR: Node of Ranvier

## References

1. Bharadwaj HM, Verhulst S, Shaheen L, Liberman MC, and Shinn-Cunningham BG. Cochlear neuropathy and the coding of supra-threshold sound. Front Syst Neurosci. 2014;8:26.

2. Gordon-Salant S. Hearing loss and aging: new research findings and clinical implications. J Rehabil Res Dev. 2005;42(4 Suppl 2):9–24.

3. Grose JH, and Mamo SK. Processing of temporal fine structure as a function of age. Ear Hear. 2010;31(6):755–60.

4. Halpin C, Thornton A, and Hasso M. Low-frequency sensorineural loss: clinical evaluation and implications for hearing aid fitting. Ear Hear. 1994;15(1):71–81.

5. Ruggles D, Bharadwaj H, and Shinn-Cunningham BG. Normal hearing is not enough to guarantee robust encoding of suprathreshold features important in everyday communication. Proc Natl Acad Sci U S A. 2011;108(37):15516–21.

6. Alvord LS. Cochlear dysfunction in “normal-hearing” patients with history of noise exposure. Ear Hear. 1983;4(5):247–50.

7. Kujala T, Shtyrov Y, Winkler I, Saher M, Tervaniemi M, Sallinen M, et al. Long-term exposure to noise impairs cortical sound processing and attention control. Psychophysiology. 2004;41(6):875–81.

8. Kumar UA, Ameenudin S, and Sangamanatha AV. Temporal and speech processing skills in normal hearing individuals exposed to occupational noise. Noise Health. 2012;14(58):100–5.

9. Liberman MC, Epstein MJ, Cleveland SS, Wang H, and Maison SF. Toward a Differential Diagnosis of Hidden Hearing Loss in Humans. PLoS One. 2016;11(9):e0162726.

10. Stone MA, Moore BC, and Greenish H. Discrimination of envelope statistics reveals evidence of sub-clinical hearing damage in a noise-exposed population with ‘normal’ hearing thresholds. Int J Audiol. 2008;47(12):737–50.

11. Pichora-Fuller MK, and Souza PE. Effects of aging on auditory processing of speech. Int J Audiol. 2003;42 Suppl 2:2S11–6.

12. Frisina DR, and Frisina RD. Speech recognition in noise and presbycusis: relations to possible neural mechanisms. Hear Res. 1997;106(1-2):95–104.

13. Dubno JR, Dirks DD, and Morgan DE. Effects of age and mild hearing loss on speech recognition in noise. J Acoust Soc Am. 1984;76(1):87–96.

14. Kujawa SG, and Liberman MC. Adding insult to injury: cochlear nerve degeneration after “temporary” noise-induced hearing loss. J Neurosci. 2009;29(45):14077–85.

15. Kujawa SG, and Liberman MC. Synaptopathy in the noise-exposed and aging cochlea: Primary neural degeneration in acquired sensorineural hearing loss. Hear Res. 2015;330(Pt B):191–9.

16. Lin HW, Furman AC, Kujawa SG, and Liberman MC. Primary neural degeneration in the Guinea pig cochlea after reversible noise-induced threshold shift. J Assoc Res Otolaryngol. 2011;12(5):605–16.

17. Liu L, Wang H, Shi L, Almuklass A, He T, Aiken S, et al. Silent damage of noise on cochlear afferent innervation in guinea pigs and the impact on temporal processing. PLoS One. 2012;7(11):e49550.

18. Lobarinas E, Spankovich C, and Le Prell CG. Evidence of “hidden hearing loss” following noise exposures that produce robust TTS and ABR wave-I amplitude reductions. Hear Res. 2017;349:155–63.

19. Shi L, Liu K, Wang H, Zhang Y, Hong Z, Wang M, et al. Noise induced reversible changes of cochlear ribbon synapses contribute to temporary hearing loss in mice. Acta Otolaryngol. 2015;135(11):1093–102.

20. Song Q, Shen P, Li X, Shi L, Liu L, Wang J, et al. Coding deficits in hidden hearing loss induced by noise: the nature and impacts. Sci Rep. 2016;6:25200.

21. Mohrle D, Ni K, Varakina K, Bing D, Lee SC, Zimmermann U, et al. Loss of auditory sensitivity from inner hair cell synaptopathy can be centrally compensated in the young but not old brain. Neurobiol Aging. 2016;44:173–84.

22. Liberman MC. Hidden Hearing Loss. Sci Am. 2015;313(2):48–53.

23. C. Kohrman D, Wan G, Cassinotti L, and Corfas G. Hidden Hearing Loss: A Disorder with Multiple Etiologies and Mechanisms. Cold Spring Harbor Perspectives in Medicine. 2020;10(1):a035493.

24. Schaette R, and McAlpine D. Tinnitus with a normal audiogram: physiological evidence for hidden hearing loss and computational model. J Neurosci. 2011;31(38):13452–7.

25. Ji L, Borges BC, Martel DT, Wu C, Liberman MC, Shore SE, et al. From hidden hearing loss to supranormal auditory processing by neurotrophin 3-mediated modulation of inner hair cell synapse density. PLoS Biol. 2024;22(6):e3002665.

26. Viana LM, O’Malley JT, Burgess BJ, Jones DD, Oliveira CA, Santos F, et al. Cochlear neuropathy in human presbycusis: Confocal analysis of hidden hearing loss in post-mortem tissue. Hear Res. 2015;327:78–88.

27. Wu PZ, Liberman LD, Bennett K, de Gruttola V, O’Malley JT, and Liberman MC. Primary Neural Degeneration in the Human Cochlea: Evidence for Hidden Hearing Loss in the Aging Ear. Neuroscience. 2019;407:8–20.

28. Wan G, and Corfas G. Transient auditory nerve demyelination as a new mechanism for hidden hearing loss. Nature Communications. 2017;8(1):14487.

29. Dyck PJ, and Lambert EH. Lower motor and primary sensory neuron diseases with peroneal muscular atrophy. II. Neurologic, genetic, and electrophysiologic findings in various neuronal degenerations. Arch Neurol. 1968;18(6):619–25.

30. Rossor AM, Polke JM, Houlden H, and Reilly MM. Clinical implications of genetic advances in Charcot-Marie-Tooth disease. Nat Rev Neurol. 2013;9(10):562–71.

31. Ramchandren S. Charcot-Marie-Tooth Disease and Other Genetic Polyneuropathies. Continuum (Minneap Minn). 2017;23(5, Peripheral Nerve and Motor Neuron Disorders):1360–77.

32. Li J, Parker B, Martyn C, Natarajan C, and Guo J. The PMP22 gene and its related diseases. Mol Neurobiol. 2013;47(2):673–98.

33. Snipes GJ, Suter U, Welcher AA, and Shooter EM. Characterization of a novel peripheral nervous system myelin protein (PMP-22/SR13). J Cell Biol. 1992;117(1):225–38.

34. Choi JE, Seok JM, Ahn J, Ji YS, Lee KM, Hong SH, et al. Hidden hearing loss in patients with Charcot-Marie-Tooth disease type 1A. Sci Rep. 2018;8(1):10335.

35. Choi JE, Seol HY, Seok JM, Hong SH, Choi BO, and Moon IJ. Psychoacoustics and neurophysiological auditory processing in patients with Charcot-Marie-Tooth disease types 1A and 2A. Eur J Neurol. 2020;27(10):2079–88.

36. Rance G, Ryan MM, Bayliss K, Gill K, O’Sullivan C, and Whitechurch M. Auditory function in children with Charcot-Marie-Tooth disease. Brain. 2012;135(Pt 5):1412–22.

37. Verhagen WI, Huygen PL, Gabreels-Festen AA, Engelhart M, van Mierlo PJ, and van Engelen BG. Sensorineural hearing impairment in patients with Pmp22 duplication, deletion, and frameshift mutations. Otol Neurotol. 2005;26(3):405–14.

38. Plack CJ, Leger A, Prendergast G, Kluk K, Guest H, and Munro KJ. Toward a Diagnostic Test for Hidden Hearing Loss. Trends Hear. 2016;20.

39. Joo IS, Ki CS, Joo SY, Huh K, and Kim JW. A novel point mutation in PMP22 gene associated with a familial case of Charcot-Marie-Tooth disease type 1A with sensorineural deafness. Neuromuscul Disord. 2004;14(5):325–8.

40. Kovach MJ, Campbell KC, Herman K, Waggoner B, Gelber D, Hughes LF, et al. Anticipation in a unique family with Charcot-Marie-Tooth syndrome and deafness: delineation of the clinical features and review of the literature. Am J Med Genet. 2002;108(4):295–303.

41. Kovach MJ, Lin JP, Boyadjiev S, Campbell K, Mazzeo L, Herman K, et al. A unique point mutation in the PMP22 gene is associated with Charcot-Marie-Tooth disease and deafness. Am J Hum Genet. 1999;64(6):1580–93.

42. Luigetti M, Zollino M, Conti G, Romano A, and Sabatelli M. Inherited neuropathies and deafness caused by a PMP22 point mutation: a case report and a review of the literature. Neurol Sci. 2013;34(9):1705–7.

43. Kabzinska D, Sinkiewicz-Darol E, Hausmanowa-Petrusewicz I, and Kochanski A. Charcot-Marie-Tooth type 1A disease caused by a novel Ser112Arg mutation in the PMP22 gene, coexisting with a slowly progressive hearing impairment. J Appl Genet. 2010;51(2):203–9.

44. Zhou R, Abbas PJ, and Assouline JG. Electrically evoked auditory brainstem response in peripherally myelin-deficient mice. Hear Res. 1995;88(1-2):98–106.

45. Zhou R, Assouline JG, Abbas PJ, Messing A, and Gantz BJ. Anatomical and physiological measures of auditory system in mice with peripheral myelin deficiency. Hear Res. 1995;88(1-2):87–97.

46. Verhamme C, King RH, ten Asbroek AL, Muddle JR, Nourallah M, Wolterman R, et al. Myelin and axon pathology in a long-term study of PMP22-overexpressing mice. J Neuropathol Exp Neurol. 2011;70(5):386–98.

47. Suter U, Moskow JJ, Welcher AA, Snipes GJ, Kosaras B, Sidman RL, et al. A leucine-to-proline mutation in the putative first transmembrane domain of the 22-kDa peripheral myelin protein in the trembler-J mouse. Proc Natl Acad Sci U S A. 1992;89(10):4382–6.

48. Budak M, Grosh K, Sasmal A, Corfas G, Zochowski M, and Booth V. Contrasting mechanisms for hidden hearing loss: Synaptopathy vs myelin defects. PLoS Comput Biol. 2021;17(1):e1008499.

49. Kobel M, Le Prell CG, Liu J, Hawks JW, and Bao J. Noise-induced cochlear synaptopathy: Past findings and future studies. Hear Res. 2017;349:148–54.

50. Anniko M. Early development and maturation of the spiral ganglion. Acta Otolaryngol. 1983;95(3-4):263–76.

51. Wang J, Zhang B, Jiang H, Zhang L, Liu D, Xiao X, et al. Myelination of the postnatal mouse cochlear nerve at the peripheral-central nervous system transitional zone. Front Pediatr. 2013;1:43.

52. Sergeyenko Y, Lall K, Liberman MC, and Kujawa SG. Age-related cochlear synaptopathy: an early-onset contributor to auditory functional decline. J Neurosci. 2013;33(34):13686–94.

53. Hickox AE, and Liberman MC. Is noise-induced cochlear neuropathy key to the generation of hyperacusis or tinnitus? J Neurophysiol. 2014;111(3):552–64.

54. Asokan MM, Williamson RS, Hancock KE, and Polley DB. Sensory overamplification in layer 5 auditory corticofugal projection neurons following cochlear nerve synaptic damage. Nat Commun. 2018;9(1):2468.

55. Resnik J, and Polley DB. Cochlear neural degeneration disrupts hearing in background noise by increasing auditory cortex internal noise. Neuron. 2021;109(6):984–96 e4.

56. Knipper M, Van Dijk P, Nunes I, Ruttiger L, and Zimmermann U. Advances in the neurobiology of hearing disorders: recent developments regarding the basis of tinnitus and hyperacusis. Prog Neurobiol. 2013;111:17–33.

57. Bai Y, Treins C, Volpi VG, Scapin C, Ferri C, Mastrangelo R, et al. Treatment with IFB-088 Improves Neuropathy in CMT1A and CMT1B Mice. Mol Neurobiol. 2022;59(7):4159–78.

58. Fledrich R, Stassart RM, Klink A, Rasch LM, Prukop T, Haag L, et al. Soluble neuregulin-1 modulates disease pathogenesis in rodent models of Charcot-Marie-Tooth disease 1A. Nat Med. 2014;20(9):1055–61.

59. Henry EW, Cowen JS, and Sidman RL. Comparison of Trembler and Trembler-J mouse phenotypes: varying severity of peripheral hypomyelination. J Neuropathol Exp Neurol. 1983;42(6):688–706.

60. Prior R, Verschoren S, Vints K, Jaspers T, Rossaert E, Klingl YE, et al. HDAC3 Inhibition Stimulates Myelination in a CMT1A Mouse Model. Mol Neurobiol. 2022;59(6):3414–30.

61. Robertson AM, King RH, Muddle JR, and Thomas PK. Abnormal Schwann cell/axon interactions in the Trembler-J mouse. J Anat. 1997;190 ( Pt 3)(Pt 3):423–32.

62. Robertson AM, Perea J, McGuigan A, King RH, Muddle JR, Gabreels-Festen AA, et al. Comparison of a new pmp22 transgenic mouse line with other mouse models and human patients with CMT1A. J Anat. 2002;200(4):377–90.

63. Hossain WA, Antic SD, Yang Y, Rasband MN, and Morest DK. Where is the spike generator of the cochlear nerve? Voltage-gated sodium channels in the mouse cochlea. J Neurosci. 2005;25(29):6857–68.

64. Kim KX, and Rutherford MA. Maturation of NaV and KV Channel Topographies in the Auditory Nerve Spike Initiator before and after Developmental Onset of Hearing Function. J Neurosci. 2016;36(7):2111–8.

65. Rutherford MA, Chapochnikov NM, and Moser T. Spike Encoding of Neurotransmitter Release Timing by Spiral Ganglion Neurons of the Cochlea. The Journal of Neuroscience. 2012;32(14):4773–89.

66. Arroyo EJ, and Scherer SS. On the molecular architecture of myelinated fibers. Histochem Cell Biol. 2000;113(1):1–18.

67. Boggs JM. Myelin basic protein: a multifunctional protein. Cell Mol Life Sci. 2006;63(17):1945–61.

68. Gabreels-Festen AA, Bolhuis PA, Hoogendijk JE, Valentijn LJ, Eshuis EJ, and Gabreels FJ. Charcot-Marie-Tooth disease type 1A: morphological phenotype of the 17p duplication versus PMP22 point mutations. Acta Neuropathol. 1995;90(6):645–9.

69. Giuliani N, Holte L, Shy M, and Grider T. The audiologic profile of patients with Charcot-Marie Tooth neuropathy can be characterised by both cochlear and neural deficits. Int J Audiol. 2019;58(12):902–12.

70. Anzalone CL, Nuhanovic S, Olund AP, and Carlson ML. Cochlear Implantation in Charcot-Marie-Tooth Disease: Case Report and Review of the Literature. Case Rep Med. 2018;2018:1760978.

71. Kobayashi M, Yoshida T, Sugimoto S, Teranishi M, Hara D, Kimata Y, et al. Cochlear implantation in patient with Charcot-Marie-Tooth disease. Auris Nasus Larynx. 2021;48(2):327–30.

72. Uchida Y, Sugiura S, Nishita Y, Saji N, Sone M, and Ueda H. Age-related hearing loss and cognitive decline - The potential mechanisms linking the two. Auris Nasus Larynx. 2019;46(1):1–9.

73. Griffiths TD, Lad M, Kumar S, Holmes E, McMurray B, Maguire EA, et al. How Can Hearing Loss Cause Dementia? Neuron. 2020;108(3):401–12.

74. Chern A, and Golub JS. Age-related Hearing Loss and Dementia. Alzheimer Dis Assoc Disord. 2019;33(3):285–90.

75. Fledrich R, Akkermann D, Schutza V, Abdelaal TA, Hermes D, Schaffner E, et al. NRG1 type I dependent autoparacrine stimulation of Schwann cells in onion bulbs of peripheral neuropathies. Nat Commun. 2019;10(1):1467.

76. Sahenk Z, and Ozes B. Gene therapy to promote regeneration in Charcot-Marie-Tooth disease. Brain research. 2020;1727:146533.

77. Belin S, Ornaghi F, Shackleford G, Wang J, Scapin C, Lopez-Anido C, et al. Neuregulin 1 type III improves peripheral nerve myelination in a mouse model of congenital hypomyelinating neuropathy. Hum Mol Genet. 2019;28(8):1260–73.

78. Pisciotta C, and Pareyson D. Gene therapy and other novel treatment approaches for Charcot-Marie-Tooth disease. Neuromuscul Disord. 2023;33(8):627–35.

79. Ozes B, Myers M, Moss K, McKinney J, Ridgley A, Chen L, et al. AAV1.NT-3 gene therapy for X-linked Charcot-Marie-Tooth neuropathy type 1. Gene Ther. 2022;29(3-4):127–37.

80. Wan G, Gomez-Casati ME, Gigliello AR, Liberman MC, and Corfas G. Neurotrophin-3 regulates ribbon synapse density in the cochlea and induces synapse regeneration after acoustic trauma. Elife. 2014;3.

