## Supplementary material for "Hidden hearing loss in a Charcot-Marie-Tooth type 1A mouse model": Suplementary Figures

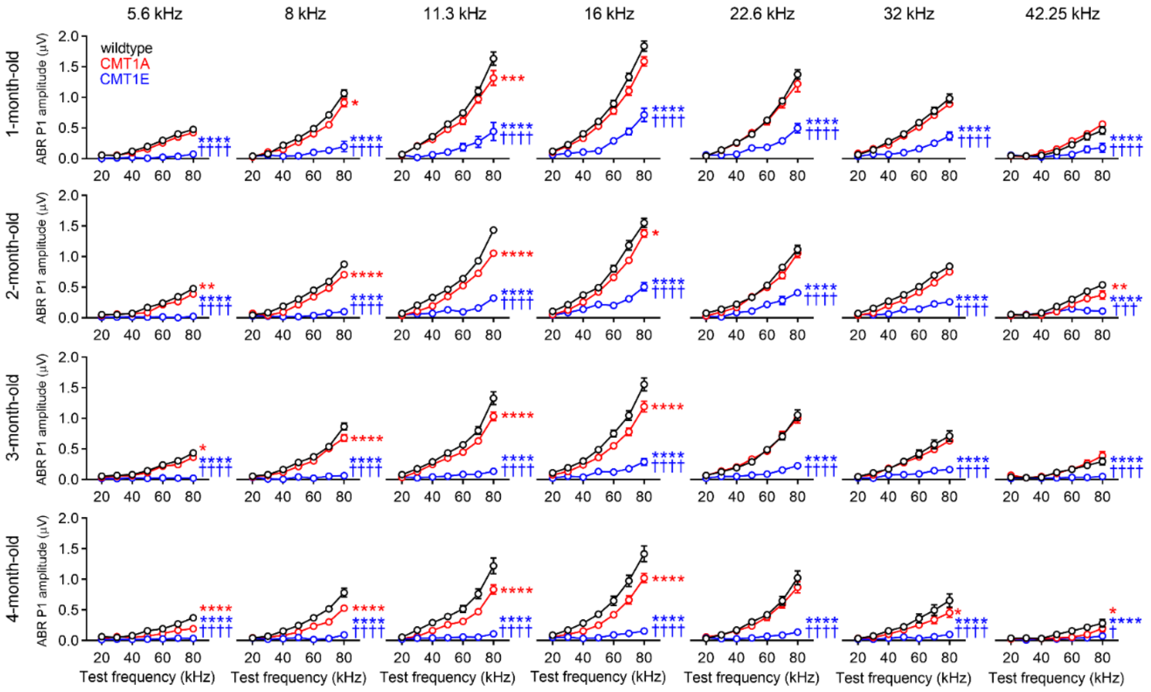


**Supplemental Figure 1. ABR input-output functions reveal difference between CMT1A and CMT1E auditory phenotypes.** ABR peak I amplitudes in CMT1A animals show progressive defects, staring at 8-11 kHz and expanding to other frequencies with aging. In contrast, all frequency areas are severely affected in CMT1E mice already at one month old. Wildtype n = 14–19 mice; CMT1A n = 8–11 mice; CMT1E n = 6–12 mice. Two-way ANOVA followed by Tukey's multiple comparisons test was used to evaluate ABR peak I amplitude statistical differences among the groups at every individual time point and tested frequency. * $p < 0.05$; ** $p < 0.01$; *** $p < 0.001$; **** $p < 0.0001$ vs wildtype mice; † $p < 0.05$; ††† $p < 0.001$; †††† $p < 0.0001$ vs CMT1A mice. Error bars represent SEM.

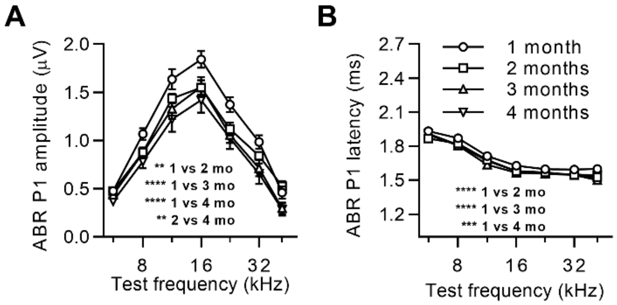


**Supplemental Figure 2. C57BL/6J wildtype mice have early onset ABR peak I amplitude decline. (A)** Wildtype mice show a small but significant progressive decline in ABR peak I amplitudes. **(B)** ABR peak I latencies shorten between 1 and 2 months old, possibly due to the maturation of AN myelination. Wildtype n = 14–19 mice. ABR peak I amplitudes and latencies were obtained at suprathreshold levels (80 dB SPL). Two-way ANOVA followed by Tukey's multiple comparisons test was used to evaluate statistical differences among the experimental groups for either ABR peak I amplitude or ABR peak I latency. ** $p < 0.01$; **** $p < 0.0001$. Error bars represent SEM.

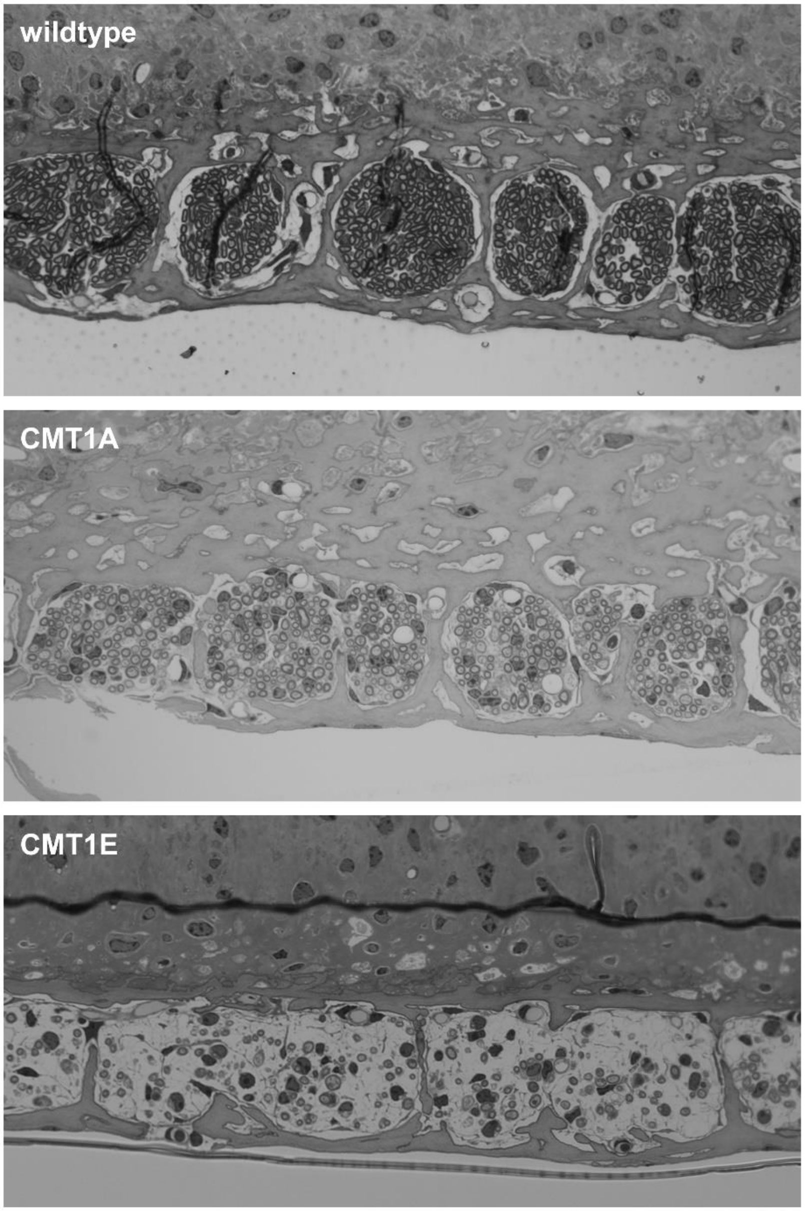


**Supplemental Figure 3.** Representative images illustrating the appearance of myelinated axon bundles in the OSL for wildtype (top), CMT1A (middle) and CMT1E (bottom) mice at 16kHz cochlear region.

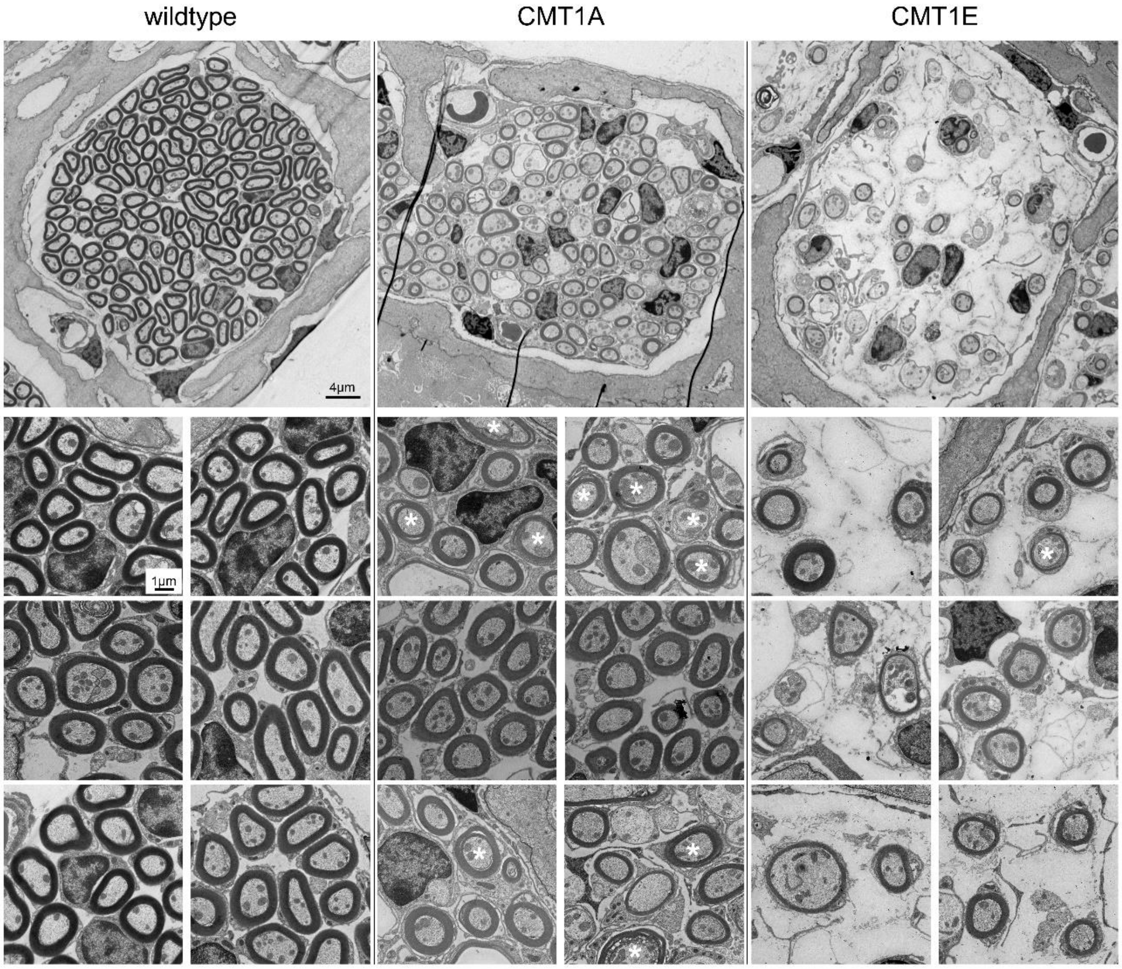


**Supplemental Figure 4.** Electron micrographs of sections through the OSL showing an axon bundle (top) and six high magnification images (two per individual mouse, top, middle, bottom, respectively; n = 3) of the axons from 4-month-old wildtype (left), CMT1A (center) and CMT1E (right) mice. Whereas some axons show myelin compaction defects in CMT1A mice (white asterisks), axonal loss and myelin abnormalities, including hypomyelination, are more dramatic in CMT1E mice.
